# Effects of Uni- and Bi-directional Interaction During Dyadic Ankle and Wrist Tracking

**DOI:** 10.1101/2024.11.25.624926

**Authors:** Matthew R. Short, Daniel Ludvig, Francesco Di Tommaso, Lorenzo Vianello, Eric J. Perreault, Emek Barış Küçüktabak, Levi Hargrove, Kevin Lynch, Etienne Burdet, Jose L. Pons

## Abstract

Haptic human-robot-human interaction allows users to feel and respond to one another’s forces while interfacing with separate robotic devices, providing customizable infrastructure for studying physical interaction during motor tasks (i.e., physical rehabilitation). For both upper- and lower-limb tasks, previous work has shown that virtual interactions with a partner can improve motor performance and enhance individual learning. However, whether the mechanism of these improvements generalizes across different human systems is an open question. In this work, we investigate the effects of haptic interaction between healthy individuals during a trajectory tracking task involving single-joint movements at the wrist and ankle. We compare tracking performance and muscle activation during haptic conditions where pairs of participants were uni- and bidirectionally connected, in order to investigate the contribution of real-time responses from a partner during the interaction. Findings indicate similar improvements in tracking performance during the bidirectional interaction for both the wrist and ankle, despite significant differences in how individuals modulated co-contraction. For each joint, bidirectional and unidirectional interaction resulted in similar improvements for the worse partner in the dyad. For the better partner, bidirectional interaction outperformed unidirectional interaction, likely due to changes in movement planning that were not observed in the unidirectional condition. While these results suggest that unidirectional interaction is sufficient for error correction of less skilled individuals during simple motor tasks, they also highlight the mutual benefits of bidirectional interaction which are consistent across the upper and lower limbs.

## I. Introduction

In daily life, humans interact through physical touch to assist and learn from one another. During physical rehabilitation, for instance, a therapist interacts with a patient in a number of ways, from applying corrective forces during dynamic movements to manually imposing a movement while stretching a stiff joint. However, systematically customizing and characterizing these interactions is difficult due to the challenge of measuring contact forces exchanged by two individuals simultaneously. To this end, robotic systems can be leveraged to study various aspects of human-human physical interaction. This is typically accomplished by rendering virtual haptic connections (e.g., spring-damper) between two robotic interfaces [1]–[9], allowing users to feel and respond to one another’s forces during motor tasks. Such systems have been used to quantify the effects of collaborative training while haptically interacting with a partner, in terms of task performance and learning as well as the muscle activation strategy of each partner.

Previous studies, involving both upper- [1], [3], [5], [6] and lower-limb [8], [9] systems, have shown that pairs of healthy individuals (i.e., dyads) perform dynamic tracking tasks better while connected compared to tracking alone. These improvements depend on the ability (i.e., relative skill level) of each partner, as well as the stiffness of the virtual connection [3], [9]. Furthermore, better partners in each dyad exert greater effort, measured as an increase in muscle activation or cocontraction, to compensate for the inferior ability of the worse partner during the interaction [5]. However, this difference in effort can be balanced without degrading tracking improvements by using a higher stiffness on the worse partner [6]. Particularly relevant to physical rehabilitation, a few studies have found that individuals can learn tracking tasks more effectively after haptic interaction with a partner, compared to training alone [1], [7] or with conventional robotic guidance toward a reference trajectory [4].

Though these benefits of human-human physical interaction have been confirmed with various robotic interfaces involving one to three degrees-of-freedom (DoF), the mechanism of these improvements is disputed. Previous work on upper-limb dyadic behaviors suggests that individuals extract motion plan information from their partner(s) using haptic feedback [2],, [10], which they combine with their own information in a stochastically optimal way. However, our work in the lower limb suggests a mechanism for these improvements based solely on the interaction mechanics [8], [9]. In these works, we simulated dyad trials by modeling the haptic interaction as three springs in series, where each participant’s simulated angle was influenced by the physiological stiffness of their own ankle, the stiffness of the virtual spring, and the physiological stiffness of their partner’s ankle. Using trajectories recorded during unconnected trials to represent each partner’s planned trajectory, simulated trajectories are effectively a weighted average of two partially correlated signals (i.e., two users following a common trajectory). A key difference in these proposed models is the consideration of the planned trajectory. In the mutual planning model [2], [3], [10], the planned movements of each partner are estimated considering the haptic information received during the interaction. In the mechanics-based model [8], [9], planned movements are assumed to be the same with or without the interaction.

Due to differences in robotic infrastructure and task constraints across the aforementioned dyadic studies, it is unclear whether a single mechanism can explain these improvements, or whether the two proposed mechanisms, mutual planning [3] and interaction mechanics [9], are specific to the upper and lower limbs, respectively. If the basis for these mechanisms is mechanical and unrelated to mutual adaptations in planning, then changes in the movement strategy of each partner should have a limited effect on the resulting behavior. In this case, one would expect similar tracking improvements whether two individuals are connected bidirectionally (i.e., two-way spring-damper), or unidirectionally (i.e., one-way spring-damper). Unidirectional interaction has been studied less extensively in dyads, but is similar to common approaches in robotic training where guidance is provided towards a reference trajectory [1], [4], [11]; in a dyadic case, partners would be unable to share information of their motor plan as only one individual receives haptic feedback. In the context of physical rehabilitation, comparing these approaches is important as it can clarify the value of two-way interaction, for instance a physical therapist receiving feedback of their patient’s movements, during partnered training.

To address these questions in the upper and lower limb, we present results comparing dyadic behaviors during a 1-DoF trajectory tracking task at the wrist (i.e., flexion and extension) and ankle (i.e., plantarflexion and dorsiflexion). Despite differences in the functional roles of each joint, previous works have suggested that velocity control of the wrist and ankle are best explained by a common kinematic model, suggesting an invariant strategy employed by the central nervous system to minimize end effector error when performing single-joint, “pointing” tasks [12], [13]. Therefore, we expect connected partners to improve similarly during wrist and ankle tasks according to their partner’s ability, with the better partner increasing their muscle activation to compensate for the worse partner’s performance. In addition, we expect that tracking improvements at the wrist and ankle can be explained by the mechanics of the interaction. This means that partners should improve similarly whether they are connected uni- or bidirectionally, based on our previous simulation work during 1-DoF tracking [8], [9] and the limited effects of haptic assistance during spatial tasks with reduced complexity [14]. Our analysis focuses on changes in tracking error to assess improvements in performance as well as muscle co-contraction to suggest changes in human mechanics as a result of the connection. For the haptic conditions, we compare these measures between bidirectional interactions where individuals are connected to their partner in real-time and unidirectional interactions where individuals are connected to a recording of their partner’s trajectory.

## II. Methods

### A. Participants

We recruited 26 healthy individuals (12 females and 14 males; 26.5 ±3.7 years) to participate in this study and paired them into sex- and age-matched dyads, separated by a maximum of 5 years. Participants gave informed consent for their participation. The study protocol (registered on clinicaltrials.gov as NCT04578665) was conducted in accordance with the Declaration of Helsinki and approved by the institutional review board of Northwestern University (STU00212684). All participants were determined to be right-handed, assessed by the preferred writing hand. All but two participants were determined to be right-footed, assessed by the preferred foot when kicking a ball.

### B. Experimental design

Pairs of participants used either their dominant wrist or ankle to perform 1-DoF movements while strapped into two commercially available robots (M1, Fourier Intelligence, Singapore; Fig. 1A). These robots are designed for 1-DoF exercises (e.g., flexion and extension) and are equipped with a sensor to measure the interaction torque between the user and robot. In our previous work [15], a custom interaction torque controller was developed to allow transparent motion (i.e., near zero interaction torque) for each robot and to render virtual haptic interactions between users interfacing with different robots. The desired interaction torque between users *i* and *j* was calculated as

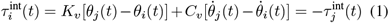

where *τ* ^int^(*t*) is the interaction torque applied to either user, *θ*(*t*) is the angular position measured by each robot, 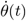 is the angular velocity, *K*_*v*_ is the virtual stiffness that is applied between the angles of the two users, and *C*_*v*_ is the virtual damping that is applied between the velocities. Motor torque commands and sensor measurements including joint angle, velocity, and interaction torque data were updated at 333 Hz.

**Fig. 1:**
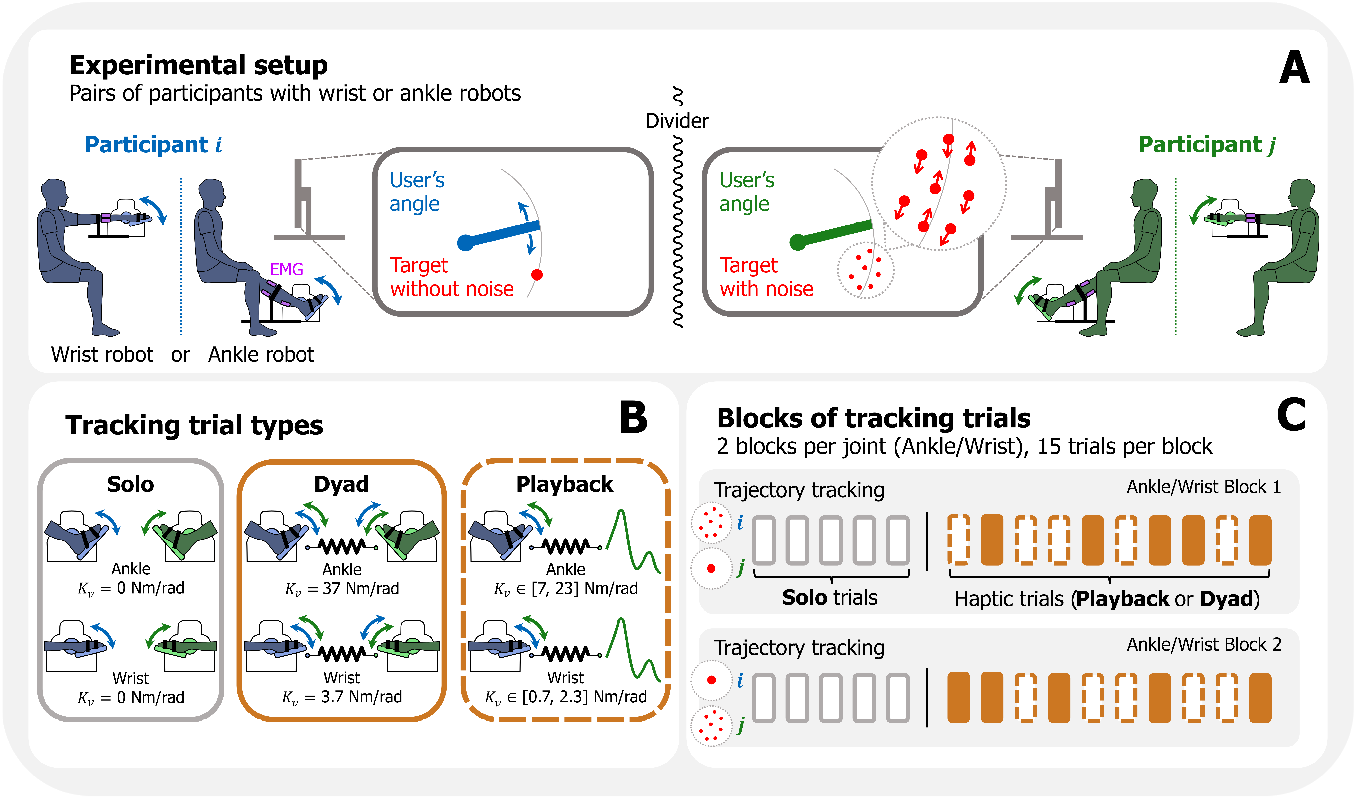
Experimental setup and block design. (A) Using either their wrists or ankles, two partners tracked multi-sine targets with different visual feedback conditions (i.e., without or with visual noise). (B) During tracking trials, three types of interaction between partners were tested: no interaction (**Solo**), bidirectional (**Dyad**) and unidirectional (**Playback**). In Dyad trials, partners interacted in real-time through compliant spring-damper elements. In Playback trials, each participant was connected to a recording of their partner’s Solo trajectory with a virtual stiffness dependent on the ability of their partner 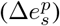,resulting in a range of stiffnesses used across participants during this condition. (C) Participants completed four blocks of tracking trials, 20 seconds in duration, for each combination of visual noise and joint (i.e., ankle or wrist); each block consisted of 5 Solo trials, followed by 10 haptic trials (Dyad or Playback in a randomized order).

To allow practical comparisons between the wrist and ankle, we made a few configurations to the robot hardware and controllers for each joint. Specifically, each robot was equipped with either a pedal for interfacing with the foot, or a manufactured handle for interfacing with the hand with the fingers splayed. The position of each participant’s foot/hand was adjusted such that their ankle/wrist joint was aligned with the robot’s center of rotation for flexion and extension movements. The interaction torque controller for the wrist and ankle configured robots was identical; manual tuning of parameters in the feedback control loop [15] was performed to achieve similar transparent tracking behaviors across joints. The experiment was divided into two phases: one for the wrist and the other for the ankle. Each phase began with an electromyography (EMG) calibration procedure followed by sets of tracking trials. The order of the wrist and ankle phases was randomized for each dyad. For the ankle experimentation, participants were seated with their knee slightly flexed and restricted to using dorsi- and plantarflexion; EMG sensors (Bagnoli, Delsys Inc., USA) were placed on the tibilias anterior (TA), gastrocnemius medialis (GM), gastrocnemius lateralis (GL) and soleus (SOL). For the wrist, participants were seated with their forearm supported and restricted to flexion and extension with the wrist in 90° of pronation; EMG sensors were placed on the extensor carpi radialis longus (ECRL) and flexor carpi radialis (FCR). A ground electrode, for both the wrist and ankle measurements, was placed on the non-dominant elbow of each participant. EMG data were collected at 333 Hz and synchronized with data from the robots using a data acquisition board (USB-6218, National Instruments, USA) and a custom Python script. Raw EMG data from the isometric calibration and tracking trials were high-pass filtered at 20 Hz (Butterworth, second order), rectified and low-pass filtered at 5 Hz (Butterworth, second order) to obtain the linear envelope of each muscle’s activation.

### C. Trajectory tracking trials

To study the effects of physical interaction during voluntary movement, we designed a continuous, dynamic tracking task involving 1-DoF. During tracking trials, dyads tried to match either their wrist or ankle angle, *θ*(*t*), to a visually-displayed, sinusoidal trajectory while their robots were commanded with the interaction torque controller described previously. *θ*(*t*) was offset, such that 0° corresponded to the center of each participant’s active range of motion. The target appeared on the display for a duration of 20 seconds and varied according to a multi-sine function:

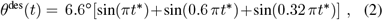

where *θ*^des^(*t*) is the instantaneous target angle presented to both participants in the dyad at a given time point (*t*). A time shift, *t*^∗^ = *t* + *t*_*r*_, with *t*_*r*_ randomly selected from a uniform distribution in the interval [0, 20] s was used to change the starting point of the trajectory for each trial and minimize memorization of the target’s movement.

To evaluate how tracking performance is affected by the ability or “skill” of each partner, we added visual noise [3] to the target of one participant in each dyad to increase their tracking error (Fig. 1A); adding visual noise in this way increases the relative differences in tracking performance between two partners. Without visual noise, the target was presented as a single 10 mm diameter point on the display. With visual noise, the target was presented as a cloud of eight 5 mm diameter points. Each point was characterized by the following randomly selected parameters: the orthogonal distance to the target *η* ∈ N(0, 10 mm), the angular distance to the target *η*_*θ*_ ∈ N(0, 4.58°), and the angular velocity 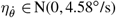.The eight points were sequentially replaced and updated with new random parameters every 250 ms.

In total, the experiment consisted of four blocks, for each combination of visual noise (i.e., *i* with noise and *j* without, *j* with noise and *i* without) and joint (i.e., wrist, ankle) (Fig. 1C). At the start of each block, participants were given time to familiarize themselves with the robot’s transparent control for the given visual condition; this involved following a multisine trajectory *θ*^des^(*t*) = 6.6°[sin(0.8 *πt*)+sin(0.4 *πt*)] for 60 seconds, followed by one tracking trial defined in Eq. (2). For the remaining trials, participants were informed that they might experience some forces from their robot, but they were blinded to the nature of these forces (i.e., the virtual connection). Within each block, dyads performed 5 **Solo** trials where the robots allowed transparent motion and 10 haptic trials (**Dyad** or **Playback**) where the robots rendered a spring-damper between the joint angles of each partner according to Eq. (1). The order of Dyad or Playback trials was randomized in the set of 10 haptic trials. The same time shifts (*t*_*r*_) selected for the Solo trials were used in the Dyad and Playback trials, such that partners were presented the same 5 trajectories across the three trial types within a block.

In the Dyad trials, the two partners received and applied haptic feedback to one another via bidirectional interaction. This interaction was achieved using the real-time joint angles of each partner and a virtual stiffness 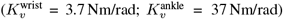 and damping 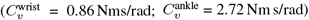. These virtual stiffnesses were selected using a similar approach to the haptic tracking experiment presented in [3]. In pilot experiments, individuals were virtually connected, using different virtual stiffnesses, to a multi-sine trajectory and asked to follow the forces they felt without the aid of visual feedback; the virtual stiffness values which produced similar tracking errors for the ankle and wrist were selected for each joint, respectively. The range of virtual stiffnesses tested for each joint during piloting was selected based on active stiffness values reported in previous works for the wrist [16] and ankle [17]. Damping constants were selected such that the system <u>wa</u>s overdamped for stability; the damping ratio 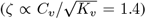 of each connection was the same.

In the Playback trials, partners received haptic feedback via unidirectional interaction. Participants were connected to a recording of their partner’s solo trajectory (featuring the same initial time shift: *t*_*r*_) with a virtual stiffness that was variable across participants 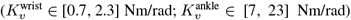 and damping consistent with the Dyad trials. Illustrated in Fig. 2, a range of virtual stiffnesses in this condition was implemented to account for the spring-series connection between the joints (i.e., human stiffness elements) of two users [9]. Partner stiffness values (*K*) were computed as a function of each partner’s ability (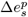, defined in section II-F), assuming that worse partners were more relaxed and better partners stiffer [5], [9]. Based on the literature values of active joint stiffness referenced previously [16], [17], we generated sigmoid functions to relate differences in Solo trial performance across pairs of participants to their expected joint stiffness during connected trials:

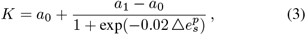

where *a*_0_, *a*_1_ are the sigmoid parameters defined separately for the wrist (*a*_0_ = 0.1 Nm/rad, *a*_1_ = 8.0 Nm/rad) and ankle (*a*_0_ = 1.0 Nm/rad, *a*_1_ = 80 Nm/rad). Sigmoid functions were used to avoid negative stiffness values of *K*. The Playback virtual stiffness was calculated by taking the series equivalent of the virtual connection stiffness in the Dyad trials and the computed partner stiffness: (1*/K*_*v*_ + 1*/K*)^−1^. Rescaling the virtual stiffness in this way accounts for the human stiffness element of each partner, which was not considered in previous work involving unidirectional interaction during dyadic tracking [1].

**Fig. 2:**
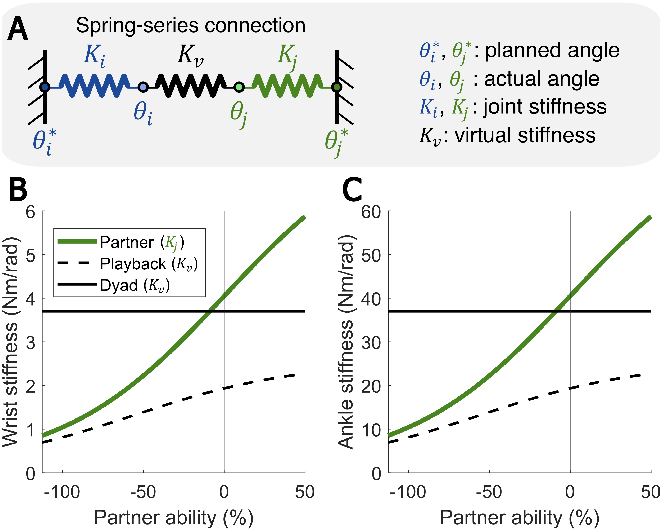
(A) Spring-series representation of the mechanical interaction between two users. Implemented values of partner joint stiffness *K*_*j*_ and virtual connection stiffness *K*_*v*_ are plotted with respect to partner ability and displayed for the (B) wrist and (C) ankle. Dyad virtual stiffness was defined as a constant value 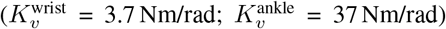 for all participants. Playback virtual stiffness was defined separately for each participant, by taking the series equivalent of the Dyad virtual stiffness and partner stiffness (1*/K*_*v*_ + 1*/K*_*j*_)^−1^, resulting in a range of stiffness values during these trials 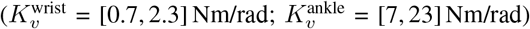.

### D. Simulating trials based on mechanics

To evaluate the contribution of mechanics during ankle and wrist tracking with a partner, we modeled the interaction as three springs in series (Fig. 2A), consistent with our previous works [8], [9], where each participant’s simulated angle was influenced by the physiological stiffness of their own joint (*K*_*i*_), the stiffness of the virtual spring (*K*_*v*_), and the physiological stiffness of their partner’s joint (*K*_*j*_). We compute the relative displacement of one spring to another, simulating the mechanical effects of the interaction on each user’s planned angle. We used each dyad’s trajectories during Solo trials (i.e., no connection) to represent their intended motion in the spring-series connection and balanced the torques across each spring:

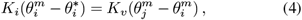

where *θ*^*m*^ is the simulated trajectory of each partner, *K* is the physiological joint stiffness of each partner, and *θ*^∗^ is each partner’s actual trajectory during solo tracking trials. Simulated trajectories were calculated by solving for *θ*^*m*^ in the following:

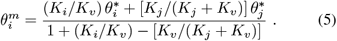

Joint stiffness values for the wrist and ankle were again calculated according to differences in partner ability 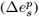 with Eq. (3).

### E. EMG calibration

EMG data from isometric torque matching trials were used to estimate torques exerted during the trajectory tracking trials. Participants performed isometric trials while fixed in the center of their active range of motion, matching a visual display of their applied torque to a series of target torque values for a period of 7 seconds (wrist: [-3, -2, -1, 1, 2, 3] Nm; ankle: [- 6, -4, -2, 2, 4, 6] Nm). Trials were repeated 2 to 3 times for each target torque with 10 seconds of rest between trials. The envelopes of EMG activity obtained during these isometric matching trials and at rest were regressed with the applied torque to generate models relating muscle activation to torque.

Two models characterized the flexor (plantarflexor) and extensor (dorsiflexor) torques (*τ*_f_(*t*), *τ*_e_(*t*)) for each joint:

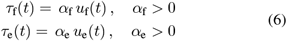

where *u*_f_(*t*), *u*_e_(*t*) are the EMG envelopes for the flexor and extensor muscles, respectively. For the wrist, the ECRL represents the extensor muscle and the FCR represents the flexor in these equations. For the ankle, TA represents the extensor muscle and one of MG, LG or SOL was selected to represent the flexor. This was decided based on the highest variance explained when using each muscle as an input for the flexor model in Eq. (6). Across all participants, this procedure resulted in appropriate fits of the isometric data (Wrist: *R*^2^ = 0.89 *±*0.09, Ankle: *R*^2^ = 0.85*±* 0.08), and was utilized to predict torque measures with EMG during dynamic tracking trials.

### G. Outcome measures

Our primary outcome measures during the trajectory tracking trials were the changes in tracking error and muscle co-contraction during the haptic conditions. All measures defined below were averaged across trials of the same condition, with respect to each joint (i.e., wrist or ankle), visual condition (i.e., with or without noise) and tracking trial type (i.e., Solo, Playback, Dyad or simulated). Each participant’s tracking error (*e*) was quantified for each trial as the root-mean-square error between the target and joint trajectories

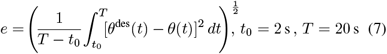

computed in the range of 2 to 20 seconds, excluding the first 2 seconds of each trial to account for initiation of task. For each joint and visual noise condition, tracking improvements were measured by taking the normalized difference between mean Solo (*e*_*s*_) and Dyad (*e*_*d*_), Playback (*e*_*pb*_) or simulated (*e*_*m*_) tracking errors. Positive values of the tracking improvements indicate better performance in the Dyad (Δ*e*_*d*_ = 100(1 −*e*_*d*_*/e*_*s*_)), Playback (Δ*e*_*pb*_ = 100(1 − *e*_*pb*_*/e*_*s*_)) or simulated trials (Δ*e*_*m*_ = 100(1 − *e*_*m*_*/e*_*s*_)) compared to Solo. Partner ability was measured by taking the normalized difference between the mean Solo tracking errors of each partner in a dyad 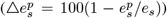). Positive values of partner ability indicate the worse partner in the pair, evaluated when performing the task alone.

Subject-specific models relating EMG activation to torque were used to quantify the mean co-contraction torque (*c*) of the agonist-antagonist pair [5]:

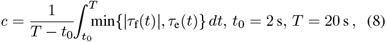

computed in the range of 2 to 20 seconds. For each participant, co-contraction torques were normalized based on the maximum (*c*_max_) and minimum (*c*_min_) values observed across all trials for each joint:

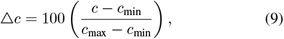

reported as mean percentages for the Dyad (Δ*c*_*d*_) and Play-back (Δ*c*_*pb*_) conditions.

### G. Statistical analysis

The goal of this study was to assess how task performance improvements and muscle activation strategies change for the wrist and ankle during uni- and bidirectional dyadic interaction. We tested two primary hypotheses for our experimental results: (1) for both the wrist and ankle, task performance improvements are different between the unidirectional and bidirectional tracking conditions, (2) improvements are different during the bidirectional condition between the wrist and ankle. As a secondary analysis, we assessed whether distinct muscle activation strategies were utilized between the wrist and ankle during haptic trials, with the expectation that better partners will exhibit higher levels of agonist-antagonist co-contraction compared to worse partners for both joints. To assess the contribution of mechanics to tracking improvements, we compare experimental results with simulated performances based on our spring-series model of the interaction.

To test our hypotheses related to changes in task performance, we used a mixed effects model with the tracking improvement in the haptic trials as a dependent variable, haptic trial type (Dyad or Playback) and joint (ankle or wrist) as categorical variables, and partner ability with linear and quadratic forms as continuous variables, as well as the interaction between each of these predictors. We used a mixed effects model with similar structure to evaluate differences in muscle activation, with normalized co-contraction torque in the haptic trials as a dependent variable, haptic trial type and joint as categorical variables, and partner ability as a continuous variable. The DoF of the mixed effects models were estimated using a Satterthwaite approximation [18]. Significance was set to 0.05 for all hypotheses related to tracking improvements and co-contraction changes. Results are presented as mean ± standard error unless otherwise specified.

## III. Results

Errors during Solo trials were similar between the ankle and wrist when tracking without visual noise (ankle: 4.1± 0.7°, wrist: 4.0± 0.8°; *t*_25_ = 0.6, *p* = 0.6) and with visual noise (ankle: 5.0 ±0.7°, wrist: 5.3± 1.0°; *t*_25_ = −1.7, *p* = 0.1). Despite this, the range of partner abilities was larger for the wrist ([− 110, 50] %) compared to the ankle ([−65, 40] %), meaning that there was a wider distribution of partner performances during wrist tracking. Improvements during the haptic trials, normalized with respect to Solo performance, increased with partner ability (Fig. 3A,B). This means that worse partners, compared to better partners, demonstrated greater improvements when haptically connected in both the Dyad and Playback trials. This relationship between partner ability and tracking improvements was characterized by a second order polynomial, including data for both the wrist and ankle during the Dyad and Playback trials (*R*^2^ = 0.79).

**Fig. 3:**
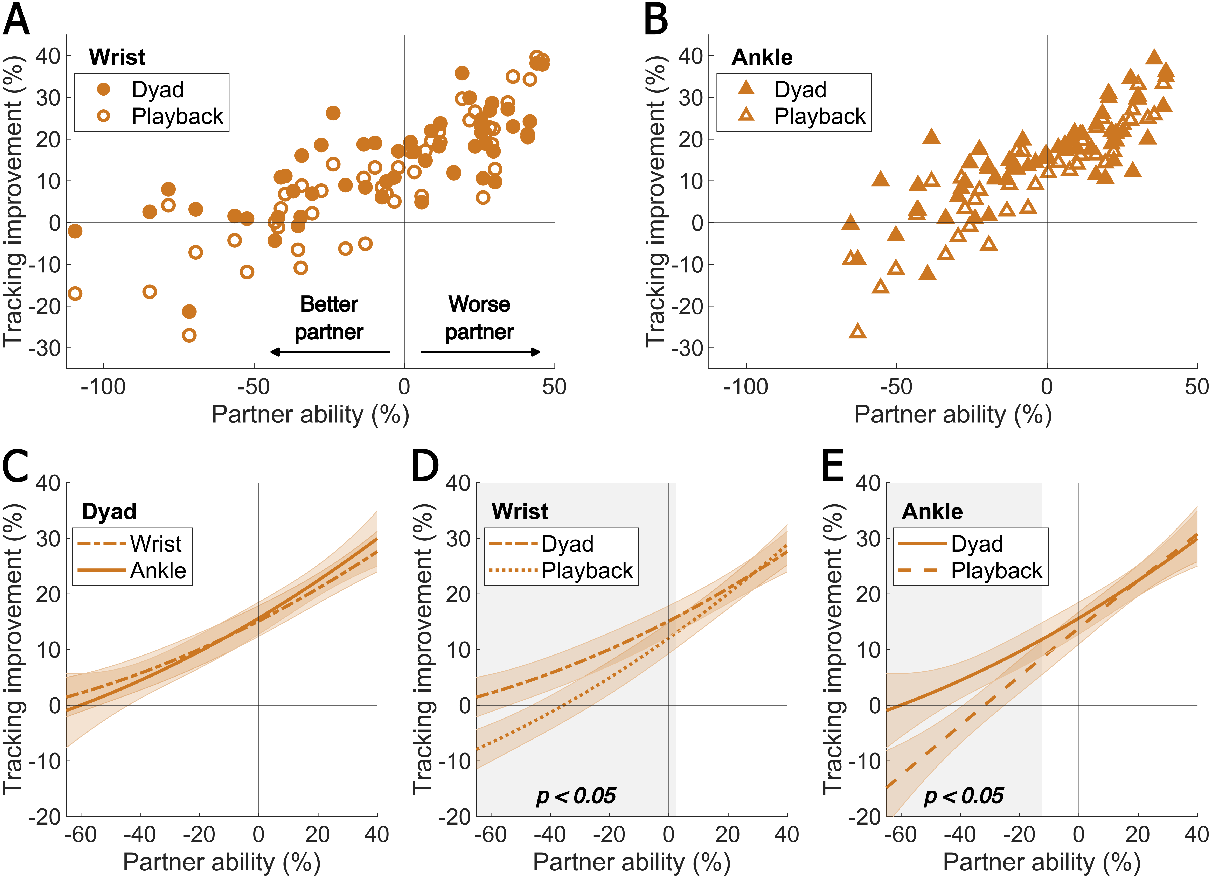
Dyadic tracking improvements were similar for the wrist and ankle, relative to each partner’s ability; better partners perform worse when connected to a recording of their partner’s trajectory (i.e., Playback), compared to a real-time connection (i.e., Dyad). Mean tracking improvements during Dyad and Playback trials plot with respect to the difference in each partner’s Solo performances are shown for the (A) wrist and (B) ankle. Linear mixed effects model fits are compared between (C) the wrist and ankle during Dyad trials, (D) Dyad and Playback trials for the wrist and (E) Dyad and Playback trials for the ankle. Vertical shaded areas indicate significant differences in the hypothesis tests that individuals improved differently as a function of partner ability for each combination of conditions.

### A. Bidirectional improvements are similar between the ankle and wrist

The improvements in tracking performance observed during the Dyad trials were similar for the wrist and ankle. Mixed effects model fits comparing ankle and wrist tracking improvements with respect to partner ability were not significantly different (*F*_3,186_ = 0.5, *p* = 0.7). Assessed over the range of partner abilities common to both joints ([−65, 40]%), there were no significant differences (*p >* 0.3) in the improvements seen during Dyad trials between the wrist and ankle (Fig. 3C). To confirm that these findings were not dependent on the 2nd order polynomial fitting of the mixed effects model, we additionally compared wrist and ankle improvements separately for better (partner ability *<* 0%) and worse (partner ability *>* 0%) partners. Again, we did not observe significant differences in wrist and ankle improvements for the better (*δ* = 1.1 *±* 1.8%, *t*_200_ = 0.4, *p* = 0.7) or worse partners (*δ* = −0.2 *±* 1.8%, *t*_200_ = −0.1, *p* = 0.9).

### B. Better partners improve less during unidirectional interaction

Better partners tracked the trajectory more accurately during the bidirectional trials (Dyad) compared to the unidirectional trials (Playback); though worse partners showed no difference in tracking performance between the two trial types. Shown in Fig. 3D,E, we observed that the tracking performance in the Playback and Dyad conditions significantly differed for both the wrist (*F*_3,184_ = 6.4, *p <* 0.001) and ankle (*F*_3,184_ = 5.6, *p <* 0.01). With the wrist, participants had significantly lower tracking improvements during Playback trials only when partner ability was below 0% (*p <* 0.05). With the ankle, participants had significantly lower tracking improvements during Playback trials when partner ability was below −16% (*p <* 0.05). Comparing improvements during Playback and Dyad trials separately for the better and worse partners, the performance of the better partners was significantly deteriorated for the wrist (*δ* = −7.2 ±1.8%, *t*_200_ = −2.9, *p <* 0.01) and ankle (*δ* = −5.9 ±1.8%, *t*_200_ = −2.4, *p <* 0.05); the performance of the worse partners during Playback and Dyad trials was not significantly different for the wrist (*δ* = 0.04 *±* 1.8%, *t*_200_ = 0.02, *p* = 1.0) or ankle (*δ* = −0.3 *±* 1.8%, *t*_200_ = −0.1, *p* = 0.9).

### C. Muscle co-contraction is modulated at the wrist, but not at the ankle, when interacting with a partner

Only at the wrist did we observe changes in muscle co-contraction dependent on partner ability during the haptic trials (Fig. 4A). Mixed effects model fits comparing ankle and wrist co-contraction during bidirectional (Dyad) trials were significantly different (*F*_2,200_ = 5.6, *p <* 0.01). At the wrist, better partners co-contracted more than worse partners during Dyad trials, indicated by a significant change in co-contraction relative to partner ability (−33.1 ±6.4%/%, *t*_200_ = −5.2, *p <* 0.001). This trend was also observed during Playback trials. Though we found no significant differences in the change co-contraction relative to partner ability (*δ* = −4.4 ±6.4%/%, *t*_200_ = −0.5, *p* = 0.6), there was a significant increase in co-contraction overall (*δ* = 7.1 ± 2.5%, *t*_200_ = 2.0, *p <* 0.05) during Playback compared to Dyad trials for the wrist.

**Fig. 4:**
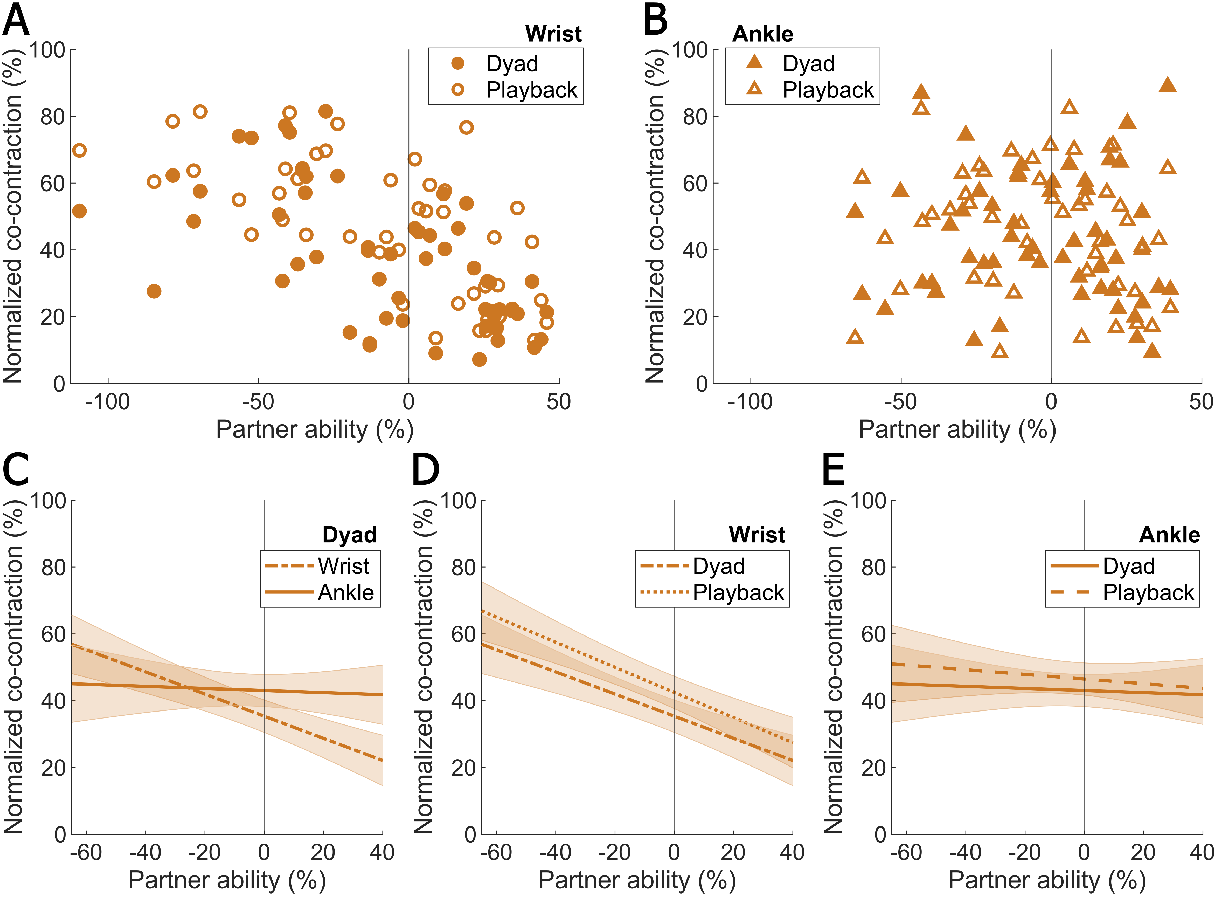
Changes in co-contraction relative to partner ability were only observed for the wrist, as better partners co-contracted more than worse partners; for the ankle, participants maintained similar levels of co-contraction regardless of their partner’s ability. Normalized co-contraction during Dyad and Playback trials plot with respect to the difference in each partner’s solo performances are shown for the (A) wrist and (B) ankle. Linear mixed effects model fits are compared between (C) the wrist and ankle during Dyad trials, (D) Dyad and Playback trials for the wrist and (E) Dyad and Playback trials for the ankle.

At the ankle, participants exhibited similar magnitudes of co-contraction independent of their partner’s ability (Fig. 4B). There was no such modulation of co-contraction in the Dyad trials, as we observed a change in co-contraction relative to partner ability that was not significantly different from zero in our mixed effects model analysis (−3.1 *±*8.7%/%, *t*_200_ = −0.4, *p* = 0.7). This was consistent for the Playback trials as well, with no significant difference in changes in co-contraction (*δ* = −3.8 *±*8.7%/%, *t*_200_ = −0.3, *p* = 0.8) between the two haptic conditions. There was slightly higher co-contraction overall during Playback trials (Fig. 4E), but this difference was not significant (*δ* = 3.4*±* 2.4%, *t*_200_ = 1.0, *p* = 0.3).

### D. Simulating improvements via mechanics underestimates bidirectional benefits

To evaluate the contribution of mechanics to improvements during the task, we simulated the interaction between partners as a system of springs in series (Fig. 2A). We created an additional model including both simulated and experimental data to compare tracking improvements with respect to partner ability across the Dyad, Playback and simulated trials; data were described by a 2nd order polynomial with partner ability (linear and quadratic forms), joint, and haptic trial type (i.e., Dyad, Playback, and simulated) as the predictors (*R*^2^ = 0.82). Shown in Fig. 5, simulated mechanical improvements underestimate the magnitude of tracking improvements during Dyad trials across the range of partner abilities for both the wrist and ankle. For the wrist, simulated improvements were quite similar to the improvements observed during the Playback trials, suggesting that mechanics were the primary mechanism of improvements during this condition. For the ankle, simulated improvements fail to characterize the improvements observed during the Playback trials. It is important to note that simulated improvements were quantified under the assumption that joint stiffness changes as a function of partner ability, which was suggested by changes in co-contraction for the wrist but not for the ankle (Fig. 4).

**Fig. 5:**
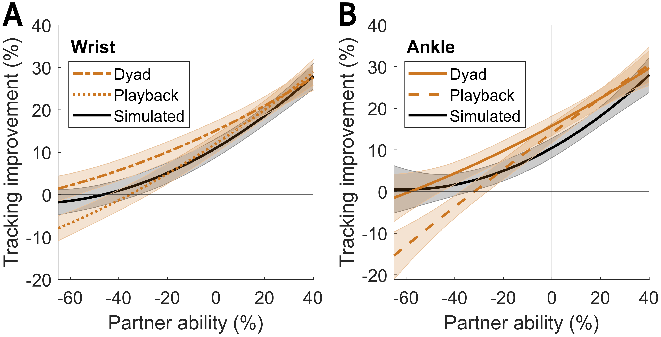
Simulated improvements based on mechanics, calculated by taking a weighted averaged of partners’ Solo trials, underestimated the improvements observed during Dyad trials. Mixed effects model fits are visualized between (A) Dyad, Playback, and **Simulated** conditions for the wrist and (B) the ankle.

## IV. Discussion

In this study, we investigated the effect of uni- and bidirectional physical interaction between healthy individuals on performance during a dynamic tracking task, in addition to a comparison of wrist and ankle behaviors during these haptic conditions. To the best of our knowledge, this was the first study to compare such behaviors across the upper and lower limbs using the same robotic infrastructure for the joints tested (i.e., wrist and ankle).

### A. Similar tracking improvements for the wrist and ankle

Comparing the wrist and ankle, we observed very similar trends in tracking improvements during real-time, bidirectional interaction with a partner. At both joints, individuals performed the 1-DoF tracking task better while connected to their partner in real-time versus alone, according to their partner’s ability. This finding is well-established and consistent with previous upper- [1], [3] and lower-limb [8], [9] studies involving compliant physical interaction between healthy individuals. However, generalization of findings across the upper and lower limbs was previously limited as these studies utilized different robots and limb postures in their experimental setups. Our work fills this gap, showing that dyadic tracking improvements can be similarly leveraged at the wrist and ankle, as long as the virtual connection stiffness is selected appropriately for the joints of interest.

A secondary, but noteworthy finding in this work was the similarities observed for the ankle and wrist during trials without interaction between partners (i.e., tracking alone). Whether trajectories were presented without or with visual noise, we found no difference in participants’ tracking errors across joints. Solo tracking errors were highly correlated between the ankle and wrist (without noise: *r* = 0.86; with noise: *r* = 0.74), meaning that the skill level of each participant generalized well across joints. Daily tasks involving these human systems are quite different, as the upper limb is typically involved in discrete, goal-directed activities like reaching and manipulation while the lower limb performs rhythmic behaviors like stepping. In contrast to the cortical-driven control of the upper limb, it is known that spinal networks are essential for locomotion in the lower limb [19], and that their function tends to be less mutable over short-term periods than cortical networks [20], [21]. Despite these distinctions, our findings are in line with previous work involving goal-directed “pointing” tasks for the ankle and wrist [12], [13], suggesting that the central nervous system uses a similar feedforward strategy to control the position of the foot or hand during ballistic movements. Our results suggest that the wrist and ankle are capable of similar performances during continuous dynamic tracking at relatively slow speeds 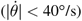 as well, though kinematic modeling of these behaviors could be tested further in future work.

### B. Unique muscle co-contraction strategies across joints

Though we observed similar tracking improvements for the wrist and ankle during bidirectional interaction, the two joints differed in terms of muscle activation during the task. Specifically, at the wrist, individuals modulated co-activation of their antagonist-agonist muscles according to the ability of their partner, perceived via the virtual connection; this was demonstrated by larger wrist co-contraction torques exerted by the better partner in the dyad compared to the worse partner. For the ankle, co-contraction torques were not dependent on partner ability and remained relatively constant for each individual throughout the tracking conditions. Despite these observed differences, our results for the ankle and wrist indicate that improvements in tracking performance can be achieved without or with changes in co-contraction of the interacting partners for these two joints, respectively.

In general, co-contraction changes in a predictable manner during the learning of novel tasks, as individuals decrease co-contraction to minimize metabolic cost and increase co-contraction to improve accuracy in response to the presented environment or task [22]. Co-contraction of the upper limb has been well-studied in the context of dynamic tasks like reaching [23], drumming [24], and trajectory tracking [5]. In the lower limb, co-contraction has primarily been studied during whole-body, loaded conditions such as walking [25] and balancing [26], and less extensively studied during isolated, dynamic tracking. Our previous work [9] showed modest changes in ankle co-contraction and overall muscle activation for better partners during dyadic tracking; however, these changes were primarily observed in a condition where individuals were rigidly connected with a very stiff spring. In the context of the compliant interaction studied in this work, it is possible that it is not advantageous or natural to modulate co-contraction with the ankle as observed at the wrist. This is supported by evidence that ankle co-contraction alone is not enough to ensure stability in unstable environments; instead, volitional descending control, associated with longer delays, is necessary to stabilize the system [27]–[29]. Therefore, our observed differences could indicate key differences in stiffness control strategies between the upper and lower limbs during dynamic tracking tasks featuring external forces.

### C. Benefits of bidirectional interaction depend on partner ability

Comparing the task performance effects of uni- and bidirectional interaction, we found that only the bidirectional interaction allowed *both* partners to improve their tracking performance, compared to tracking alone, across the full range of partner abilities. This finding was observed for both the ankle and the wrist, as better partners experienced greater tracking improvements when connected to their partner in real-time compared to a recording of their partner’s trajectory. It is important to note, however, that the worse partner in the dyad improved similarly whether or not they were connected to their partner in real-time. This differs from the results of Ganesh et al. [1], which reported that both better and worse partners tracked targets less accurately when connected to a pre-recorded trajectory during a 2-DoF upper-limb task. Our findings can be attributed to our rescaling of the virtual stiffness during the unidirectional connection, emphasizing the importance of considering the human stiffness elements involved in the interaction and the contribution of mechanics to dyadic performance.

Similarities between the worse partners’ uni- and bidirectional tracking improvements suggest that, for less skilled individuals, simply being connected to a reference trajectory which is closer to the target trajectory (i.e., the better partner’s trajectory) is sufficient for improving task performance during training; this strategy of providing assistance towards a predefined reference is often implemented in rehabilitation robotics to demonstrate desired movements to impaired individuals [30], but its effectiveness in enhancing motor learning can be limited depending on the task constraints and the skill level of the individual [31]. As our study focused on task performance effects rather than changes in individual learning, it is an open question whether interaction with a reactive partner, interaction with a human partner’s trajectory, or interaction with a “perfect” reference trajectory is the most appropriate strategy for training in healthy individuals or populations with sensorimotor impairments.

### D. Contribution of mechanics and mutual planning

To identify the contribution of mechanics to the improvements during the task, we simulated the interaction as a spring-series connection between two users’ joints [8], [9], considering the human stiffness elements (i.e., wrist or ankle stiffness) of the partners and the stiffness of the virtual connection. In this simulation, we did not consider any changes in movement planning during the interaction; any observed improvements would be attributed to mechanical error averaging between the executed movements of two users (i.e., solo, unconnected trajectories). We found that improvements estimated from this mechanical model underestimated the magnitude of improvements observed during bidirectional interaction, across the range of partner abilities. While the trend of these improvements was relatively similar between simulation and experimental results, particularly for the wrist, this shows that mechanics are likely not sufficient to fully describe the improvements observed during real-time interaction with a partner.

Though mechanics may contribute to tracking improvements during dyadic interaction, it is likely partners additionally adapt their movement plans to maximally benefit from the interaction, as suggested by previous dyadic works [2], [3]. Future work could develop a model which considers both changes in mechanics and movement planning to characterize the relative contribution of each component during dyadic interaction. In our previous work with the ankle [9], we observed closer agreement between our simulated mechanical improvements and experimental results during dyadic tracking. However, the visual feedback provided to participants was different with respect to this current work; in the previous work, we presented the same visual feedback (i.e., with or without visual noise) to both participants in the dyad, as opposed to distinct visual feedback to each partner. Providing distinct feedback in this way results in larger differences in tracking performance between partners and could potentially elicit some movement adaptation for either partner in the dyad that is not observed when partners share the same visual uncertainty of the target’s movement.

It is important to emphasize that the mechanical model evaluated in this work is limited as we do not account for real-time changes in movement planning during tracking (i.e., a user’s online prediction of the target movement) as in previous works [2], [3]. We chose this approach to isolate the effects of mechanics, independent of changes in planning, as an extension of our previous simulation [9]. Our mechanical model relies on the assumption that planned movements during the dyadic interaction are equivalent to executed movements without the interaction, which are known a priori for a given target trajectory. However, this model cannot be implemented in the case of tracking a randomly moving target where the user’s planned trajectory is unknown, thus limiting its applicability in the implementation of a more generalized and realistic robot controller [4].

### E. Limitations

One limitation of this study is the selection of partner stiffness values used in our simulation as well as the rescaling of the virtual stiffness during the unidirectional condition. Though we assumed that partner stiffness would change as a function of partner ability, this was only suggested by the trends in wrist co-contraction, but not for the ankle. Co-contraction can give an indication of how the active component of joint stiffness may change during the task, however, it is not a direct measure of joint stiffness. Furthermore, the range of partner stiffness values used in our simulation and experimentation was based on isometric data published previously for the wrist and ankle [16], [17] and it is known that stiffness of human joints can decrease considerably from postural tasks to dynamic movements [32]. Because we defined the partner stiffness values based on our hypothesis that stiffness would change as a function of partner ability, rather than attempting to estimate the stiffness during tracking, this may have affected the perceived virtual environment when participants experienced the unidirectional condition compared to the bidirectional interaction.

To address this limitation and appropriately quantify the joint stiffness of interacting users, wrist and ankle stiffness should be estimated using kinematic responses to torque perturbations during the tracking task [33]. This estimation method would also provide a continuous measure of joint stiffness, which would allow analysis of changes in mechanics within the trajectory tracking trials during different haptic conditions. However, this approach is not straightforward, as a large quantity of repeatable movements is necessary to produce accurate estimates of joint impedance [34]. In the context of our dynamic tracking task, this presents a challenge as the target trajectories are designed to appear “random” to prevent memorization of the target’s movement, resulting in a few overlapping segments across multiple trials of the same haptic conditions. In addition, external torque perturbations, which are used to estimate stiffness, must be designed in a way that they do not interfere with the torque feedback transmitted between partners during the interaction.

## V. Conclusion

During a 1-DoF trajectory tracking task, we found that bidirectional physical interaction between healthy individuals coupled at the wrist and ankle, respectively, results in similar trends of tracking improvements across joints. This was observed despite distinct muscle activation strategies across joints, as participants connected at the wrist co-contracted more when paired with a worse partner while participants connected at the ankle did not modulate co-contraction. In addition, we found that worse partners improved similarly during uni- and bidirectional interaction, while better partners were negatively affected by the unidirectional connection to the worse partner. Together, these results suggest that partners leverage both the mechanics of the virtual interaction, as well as adapt their movement plan to benefit additionally while interacting in real-time. Similarities in the response to uni- and bidirectional connections across the upper and lower limbs demonstrate that the benefits of human-human physical interaction generalize across these human systems for simple 1-DoF tracking tasks. As this work was tested in healthy individuals, future work should explore these behaviors in populations with sensorimotor impairments to determine the functional benefits of interacting with a partner during rehabilitation compared to conventional approaches in robotic assistance.

